# Chromosome-level genome assembly of *Acanthopagrus latus* using PacBio and Hi-C technologies

**DOI:** 10.1101/2021.04.08.438412

**Authors:** Dong Gao, Wenyu Fang, Ying Sims, Joanna Collins, James Torrance, Genmei Lin, Jingui Xie, Kerstin Howe, Jianguo Lu

## Abstract

The yellowfin seabream, *Acanthopagrus latus*, is widely distributed throughout the Indo-West Pacific. This fish is an ideal model species in which to study the mechanism of sex reversal since it exhibits a specific feature: sequential hermaphrodite. Here, we report a chromosome-scale assembly of the *A. latus* based on PacBio and Hi-C data. 22,485 protein-coding genes were annotated in whole genome level using transcriptome data. Taken together, this highly accurate, chromosome-level reference genome can provide a valuable resource to elucidate the mechanism of sex reversal for *A. latus*.

**Background & Summary:** Evolution of sex, especially the evolution of different sexual systems, is a fascinating subject in evolutionary biology. The Sparidae, commonly known as seabreams or progies, is a family of fishes of the order Perciformes. And this family consist about 150 species in the world, which are mainly coastal fish^1^. Previous researchers mentioned that Sparidae is an ideal taxon to study the evolution history and adaptive significance of sexual systems, particularly for both types of sequential hermaphroditism, given that this group contains many protogyny, protandry and genochorist species^2^.

The yellowfin seabream, *Acanthopagrus latus* is a protandry species which belongs to the Sparidae family. It is widely distributed in Indo-West Pacific area^3^. It has a great relevance for marine aquaculture and its biology is well focused on reproductive physiology and nutrition^4^. Interestingly, *A. latus* has a special gender feature is that it belongs to protandrous sexual system (initially as male and change later to female)^5^. Most of the past studies of *A. latus* mainly focused on the reproductive biology, population structure, aquaculture and taxonomy^3,4,6–8^. Although some sex reversal related genes were found in *A. latus*, the lack of genomic resources still limit us to elucidate the mechanism of sex reversal for this species.^9,10^. In addition, this lack was also limited the studies of evolution of sexual systems for Sparidae.

In this study, long-read (PacBio SMRT) sequencing and Hi-C sequencing technologies were applied to construct a high quality reference genome for yellowfin seabream. This high-quality genome can provide a valuable resource to elucidate the mechanism of sex reversal for *A. latus*. Furthermore, this genome can also facilitate the studies of evolution of sexual systems for Sparidae.

## Methods

### Ethics statement

All experimental procedures in our study with *A. latus* were approved by the Ethics Committee of Sun Yat-sen University.

### Sample collection, library construction and sequencing

A wild healthy female yellowfin seabream was captured from the area of Xiangzhou Bay, Zhuhai, Guangdong Province, China (Fig. 1). Seven tissues were collected respectively for genome sequencing and genome annotation, including brain, heart, liver, spleen, kidney, gonad and muscle. These samples were immediately frozen using liquid nitrogen for 30 minutes and then stored at −80°C for later usage. For high-molecular-weight (HMW) genomic DNA (gDNA) extraction, frozen samples were lysed in SDS digestion buffer with proteinase K. Then, the lysates were purified using AMPure XP beads to obtain HMW gDNA. Meanwhile, normal-molecular-weight (NMW) gDNA was extracted from the same samples using the DNeasy Blood and Tissue Kit (Qiagen, Valencia, CA, USA).

**Fig. 1.**
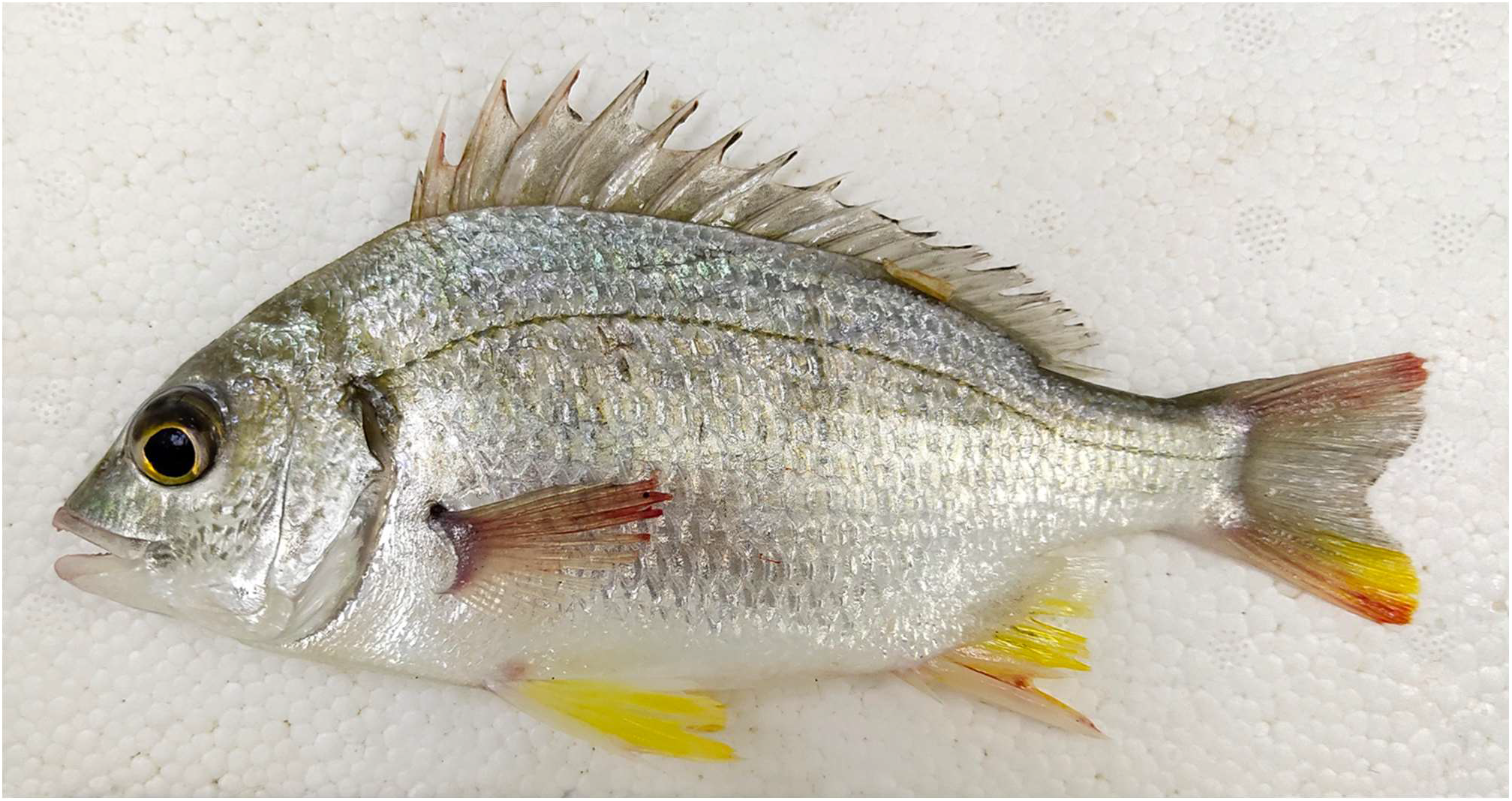
A healthy female yellowfin seabream was collected.

A whole-genome shotgun sequencing strategy was performed for genome size estimation and polishing of preliminary contigs. A library with 350 bp insert size was constructed from NMW gDNA using the standard protocol provided by Illumina (San Diego, CA, USA). Paired-end sequencing was employed using the Illumina NovaSeq platform with a read length of 2 × 150 bp. 38.44 Gbp raw reads were generated. Adaptor, low-quality and duplicated reads were trimmed with fastp (v0.20.0)^11^. In total, 34.97 Gbp clean reads were used to genome size estimation and preliminary contig polishing (Table 1).

**Table 1.**
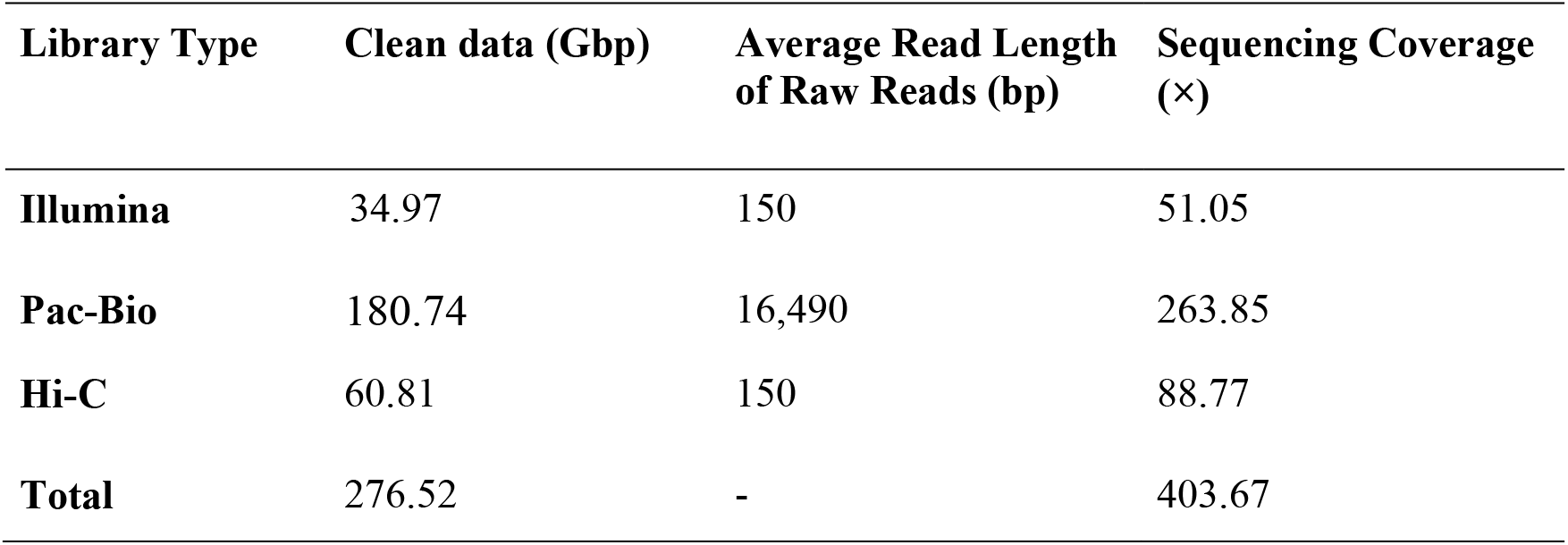
Summary of obtained data using multiple sequencing technologies. Note: The genome size of *A. latus* used to calculate sequencing coverage was 656.61 Mbp, which was estimated using a K-mer analysis of the short reads.

HWM gDNA was used in DNA template preparation for sequencing on the PacBio System following the “Template Preparation and Sequencing Guide” provided by Pacific Biosciences (Menlo Park, CA, USA). The main steps were as follows: extracted DNA was first sheared into large fragments (10 Kbp on average) and then purified and concentrated using AMPure PB beads; DNA damage and ends induced in the shearing step were repaired; blunt hairpins were subsequently ligated to the repaired fragment ends; prior to sequencing, the primer was annealed to the SMRTbell template, and then, DNA polymerase was bound to the annealed templates; finally, DNA sequencing polymerases were bound to the primer-annealed SMRTbell templates.

After sequencing, a total of 180.74 Gbp long reads were generated from the PacBio SEQUEL platform. The average length of reads was 16.49 Kbp. The long reads covered the genome about 263.85 × (Table 1).

Hi-C sequencing was performed parallel to the PacBio sequencing. We used formaldehyde to fix the conformation of the HMW gDNA. Then, the fixed DNA was sheared with DpnII restriction enzyme. The 5’ overhangs induced in the shearing step were repaired using biotinylated residues. Following the ligation of blunt-end fragments *in situ*, the isolated DNA was reverse-crosslinked, purified, and filtered to remove biotin-containing fragments. Subsequently, DNA fragment end repair, adaptor ligation, and polymerase chain reaction (PCR) were performed successively. Sequencing was performed on the Illumina NovaSeq platform and yielded a total of 67.99 Gbp paired-end reads, with an average sequencing coverage of 88.77 × (Table 1).

In addition, total RNA was extracted from each tissue using TRIZOL (Invitrogen, Carlsbad, CA, USA). The RNA samples were then treated by Dnase I. The integrity and size distribution were checked with Bioanalyzer 2100 (Agilent technologies, santa Clara, CA, USA). The high-quality RNA samples were mixed and then sequenced on Illumina NovaSeq platforms with the manufacturer’s instructions. At last, 69.36 Gbp raw reads were generated for transcriptome-base gene prediction.

### *De novo* assembly of the *A. latus* genome

In summary, whole-genome shotgun sequencing data were used in estimation of genome size and polishing of preliminary contigs; PacBio sequencing data were used for preliminary contain assembly; and Hi-C reads were used in chromosome-level scaffolding.

The shotgun sequencing data were used to estimate the genome size with Jellyfish (v2.1.3)^12^. As a result, the genome size of *A. latus* was estimated to be approximately 656.61 Mbp. All raw long-read sequences were aligned to each other using ‘dalinger’ executed by the main script of the FALCON assembler^13^. The overlap data and raw subheads were then processed to generate consensus sequences. After the error-correction step, FALCON identified the overlaps between all pairs of the preassembled error-corrected reads. The read overlaps were used to construct a directed string graph that contains sites of ‘haplotype-fused’ contigs as well as bubbles representing divergent regions between homologous sequences. Next, FALOCN-Unzip identified read haplotypes using phasing information from heterozygous positions. Phased reads were them used to assemble haplotigs and primary contigs. The shotgun sequencing data and PacBio long-reads were used to polish the preliminary contigs with Nextpolish and Arrow (version 1.21, Pacific BioSciences). The draft genome of *A. latus* was assembled in 215 contigs. The genome size equivalents to 685.14 Mbp with contig N50 of 14.88 Mbp (Table 2).

**Table 2.** Summary of the *A. latus* genome assembly and structural annotation.

| Genome Assembly |  |
| --- | --- |
| Contig N50 length (Mbp) | 14.88 |
| Number of contigs longer than N50 | 18 |
| Number of contigs | 215 |
| Scaffold N50 length (Mbp) | 30.72 |
| Number of contigs longer than N50 | 11 |
| Number of contigs | 65 |
| Total contain length (Mbp) | 685.14 |
| Structural Annotation |  |
| Number of protein-coding genes | 29,227 |
| Average genes length (bp) | 10,797.36 |
| Average exons per gene | 8.51 |
| Average exons length (bp) | 172.77 |
| Average CDS length (bp) | 1,470.37 |
| Average intro length (bp) | 1, 087.79 |

To obtain chromosome-level scaffolds, Hi-C reads were filtered in the same way as the shotgun sequencing reads. Subsequently, filtered Hi-C reads were mapped to *de novo* assembled contigs to construct contacts among the contigs using BWA (version 0.7.17)^14^with the default parameters. BAM files containing Hi-C linking messages were processed by another round of filtering, in which reads were removed if they were not mapped to the reference genome within 500 bp from the nearest restriction enzyme site. Then, LACHESIS^15^ was used for ultra-long-range scaffolding of de novo genome assemblies using the signal of genomic proximity provided by the Hi-C data (Fig. 2).

**Fig. 2.**
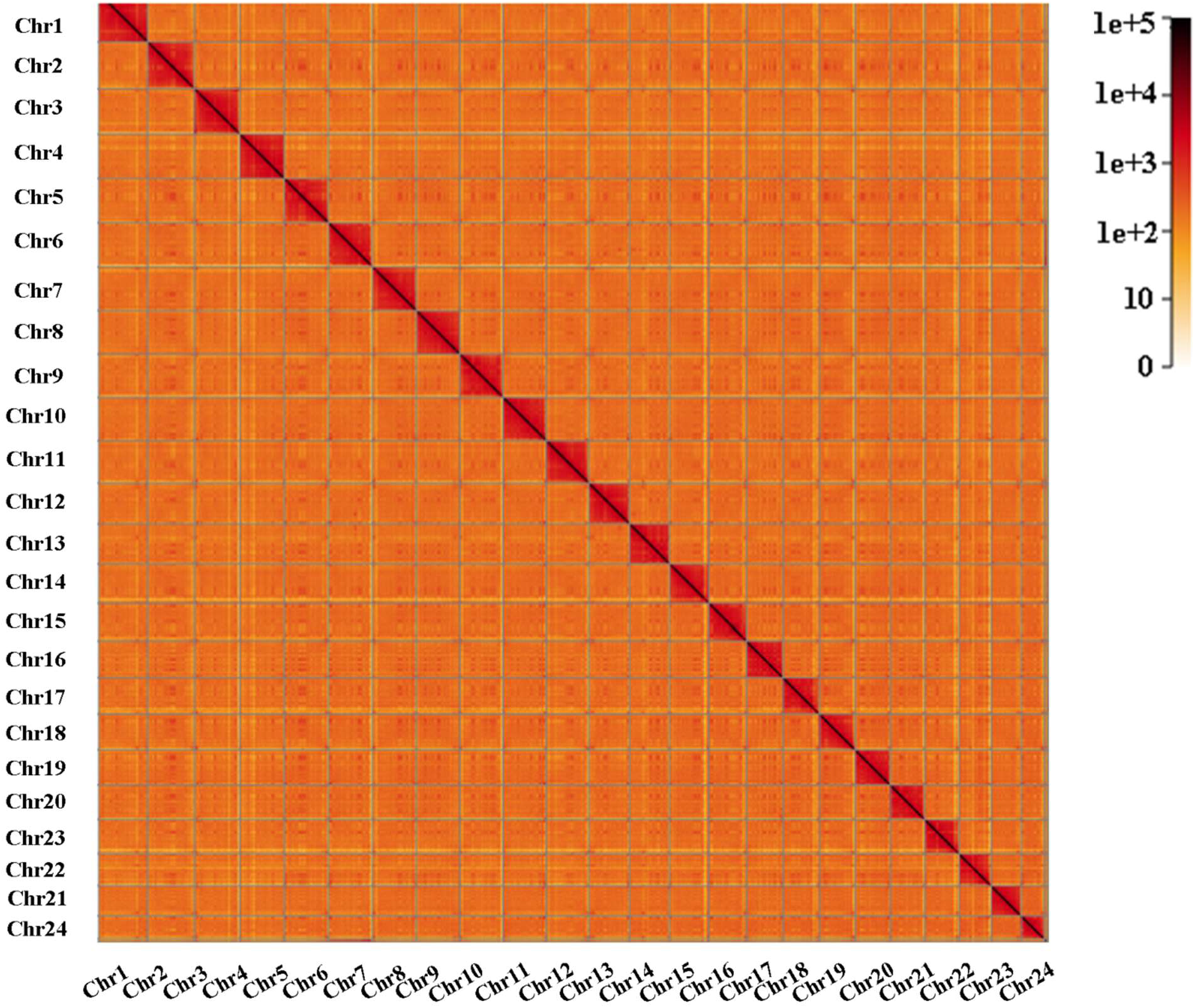
Genome-wide all by all Hi-C interation of *A. latus*.

The parameter CLUSTER_N was used to specify the number of chromosomes. For yellowfin seabream, this number was determined to be 24 in previous studies^16^. Ultimately, we obtained 24 chromosome-level scaffolds with length of 680.74 Mbp (99.36 % of the total length of genome) (Fig. 2 and Table 3).

**Table 3.** Detailed results of chromosome-level scaffolding using Hi-C technology.

| Chromosomes | Length (Mbp) | Number of Scaffolds | GC% |
| --- | --- | --- | --- |
| Chr1 | 33.53 | 1 | 41.8 |
| Chr2 | 32.02 | 1 | 42.1 |
| Chr3 | 26.36 | 1 | 42.4 |
| Chr4 | 35.91 | 1 | 41.7 |
| Chr5 | 31.40 | 1 | 41.9 |
| Chr6 | 34.04 | 1 | 41.8 |
| Chr7 | 31.19 | 1 | 42.2 |
| Chr8 | 31.34 | 1 | 42.1 |
| Chr9 | 30.72 | 1 | 41.8 |
| Chr10 | 22.41 | 1 | 42.3 |
| Chr11 | 31.32 | 1 | 41.9 |
| Chr12 | 26.68 | 1 | 41.9 |
| Chr13 | 28.11 | 1 | 42 |
| Chr14 | 22.16 | 1 | 42.7 |
| Chr15 | 28.96 | 1 | 42.1 |
| Chr16 | 25.62 | 1 | 42.6 |
| Chr17 | 31.84 | 1 | 42 |
| Chr18 | 31.74 | 1 | 41.8 |
| Chr19 | 24.96 | 1 | 42.3 |
| Chr20 | 24.49 | 1 | 42.3 |
| Chr21 | 28.39 | 1 | 42 |
| Chr22 | 26.17 | 1 | 42.1 |
| Chr23 | 25.35 | 1 | 42.6 |
| Chr24 | 16.04 | 1 | 43 |
| Total | 680.74 | 24 | - |

### Gene annotation

To obtain a fully annotated *A. latus* genome, three different approaches were employed to predicted protein-coding gens. *Ab intio* gene prediction was performed on the repeat-masked *A. latus* genome assembly using Augustus (version 3.3.1)^17^ and GeneMark-ES (version 4)^18^. Furthermore, homology-based prediction was performed using protein sequences of two common model species (*Danio rerio* and *Nile tilapia*) and two related species (*Sparus aurata* and *Larimichthys corcea*). Subsequently, these protein sequences were mapped onto the generated assembly using GeMoMa (version 1.6.1)^19^. In addition, transcriptome-based prediction was also applied by RNA-seq data. The RNA-seq reads were mapped onto the genome assembly using STAR (version 2.7.3a)^20^, and the structures of all transcribed genes were predicted by Stringtie (version 1.3.4d)^21^ with the default parameters. Based on the results of Stringtie, PASA (version 2.3.3)^22^ was used to predicted genes of genome of *A. latus*. The predicted gene sets, generated from these three approaches, were integrated to produce a non-redundant gene set using EvidenceModeler (version 1.1.1)^23^. As a result, a total of 29,227 protein-coding genes were predicted. The average number of eons per gene was 8.51. The average CDS length was 1,470.37 bp.

Gene function annotations were conducted against the NCBI nr and SwissProt protein databases, and homologs were called with E values of < 1 × 10^−5^. The functional classification of Gene Ontology (GO) categories was performed using the InterProScan program (version 5.3.2)^24^. Blastp (version 2.7.1)^25^ were performed to Kyoto Encyclopedia of Genes and Genomes (KEGG) pathway and Eukaryotic Orthologous Groups of protein (KOG) annotation analysis. As a result, a total of 22,485 genes were annotated (Fig. 3 and Table 2).

**Fig.3.**
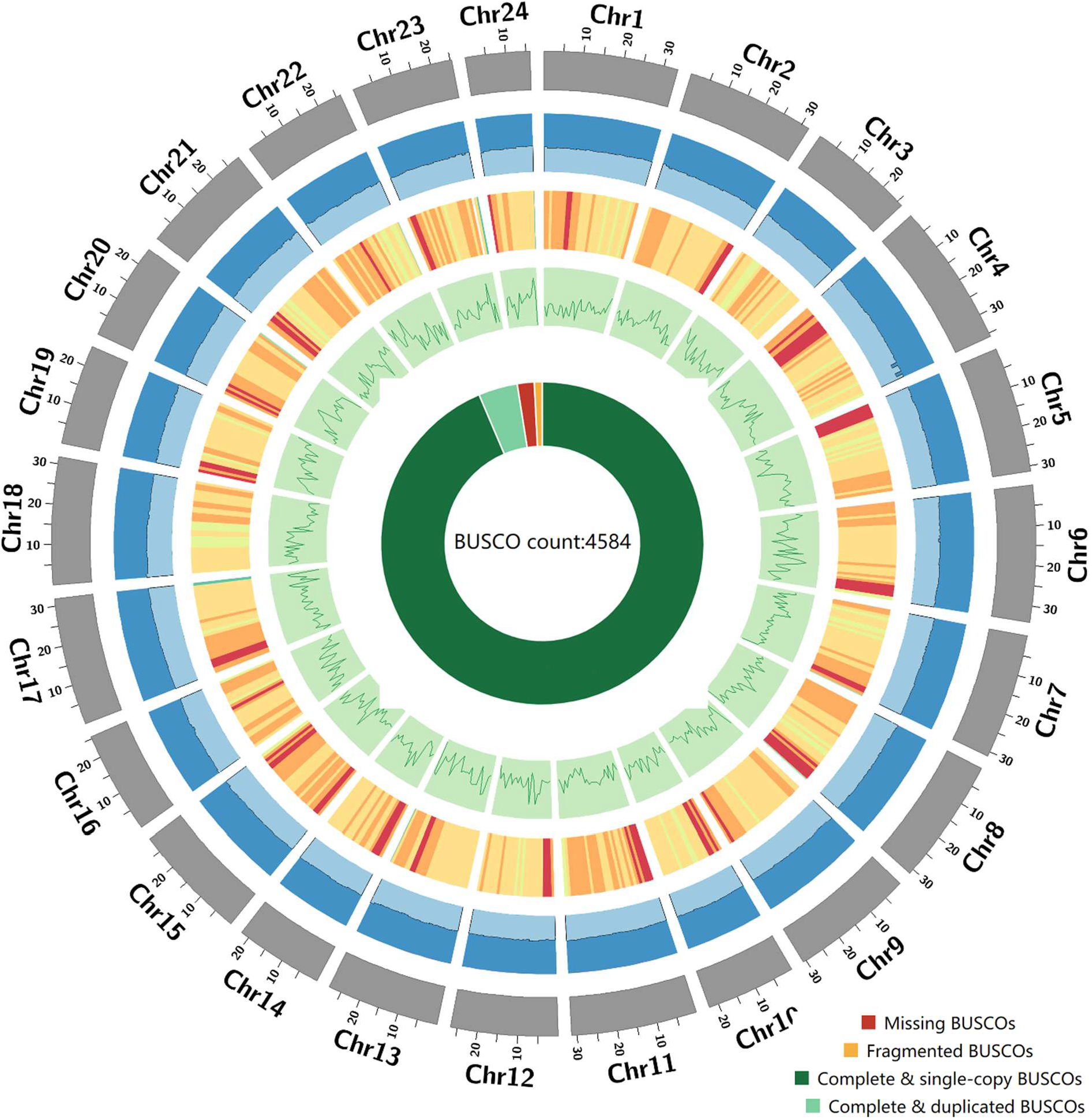
Circos plot showing 24 chromosomes of *A. latus.* Note: chromosome length in Mb unit; the blue histogram presents the GC content for 1 Mbp window; heatmap of repeats density within 1 Mbp window, ranging from 27 to 2,198 repeat sequences per million base pairs; line plot of gene density for 1 Mbp windows and the innermost pie chart displays the completeness validation results of genome assembly using BUSCO.

### Repetitive element characterization

Two software: GMATA (version 2.2)^26^ and Tandem Repeats Finder (version 4.07b)^27^ were employed to detect tandem repeats (TRs) in assembly of yellowfin seabream. In addition, tandem repeats were masked before searching transposable elements (TEs) to avoid the conflicts between TRs and TEs. A MITE database was constructed, based on masked TR genome, using MITE-hunter^28^. Meanwhile, a long terminal repeat (LTR) database was obtained by LTR_FINDER^29^ and LTRharvest^30^. Next, these two databases were combined into a TE library (TE.lib) and repeat sequences of genome were marked again. A *de novo* repeat sequence library (RepMod.lib) was generated with RepeatModeler^31^ and TEclass ^32^. TE.lib, RepMod.lib and Repbase were then integrated as a non-redundant repeat sequence library (nrRep.lib). Subsequently, *A. latus* genome was annotated with nrRep.lib using RepeatMasker (version 1.331, http://repeatmasker.org) to search repeat sequences.

Combining the annotation results of TRs and TEs, ~21.24% sequences of the *A. latus* genome were identified as repetitive elements, including 0.51% long terminal repeats (LTRs), 1.68% long interspersed nuclear elements (LINEs), 0.1% short interspersed nuclear elements (SINEs) and 5.49% of DNA transposons, (Fig. 3 and Table 4).

**Table 4.** Detailed classification of repeat sequences. Note: “Unknown” represents transposable elements that could not be classified by RepeatMasker.

| Type | Number of elements | Length of sequence (bp) | Percentage of sequence (%) |
| --- | --- | --- | --- |
| LTR | 8,942 | 3,512,788 | 0.51 |
| LINE | 44,400 | 11,480,725 | 1.68 |
| SINE | 6,094 | 686,582 | 0.10 |
| DNA | 206,827 | 37,633,093 | 5.49 |
| Tandem Repeats | 188,984 | 7,124,823 | 1.04 |
| Unknown | 508,488 | 81,118,285 | 11.84 |
| Other | 17,306 | 3,984,546 | 0.58 |
| Total Repeats | 981,041 | 145,540,842 | 21.24 |

### Data Records

All sequencing data, including Illumina short reads, PacBio long reads, Hi-C reads were submitted to the NCBI Sequence Read Archive (SRA) database under Bioproject accession PRJEB40702. The assembled genome was deposited at DDBJ/ENA/GenBank under the accession GCA_904848185.1.

### Technical Validation

The completeness and accuracy of the assembly further assessed in multiple ways. First, the Illunima reads were re-mapped onto the assembly using BWA. As a result, 99.57% of the reads were accurately mapped with a coverage of 99.70%. Subsequently, Benchmarking Universal Single-Copy Orthologs (BUSCO) software (version 3.0.1)^33^ was executed using actinopterygii_odb9 database to assess the predicted gene set. The result showed that 98.3% of all 4584 BUSCOs were assembled, including 97.50% and 0.80% of all BUSCOs were completely and partially assembled, also implying a high level of completeness for the *de novo* assembly (Table 5).

**Table 5.**
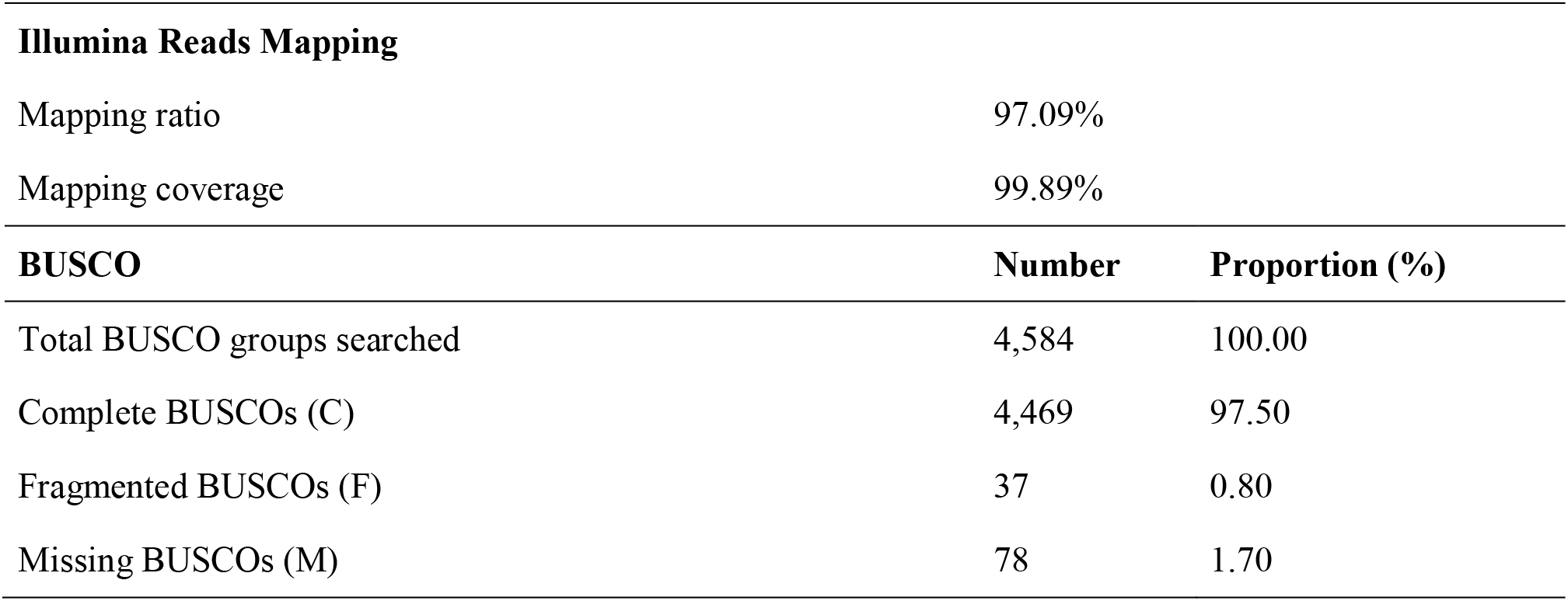
Details of accuracy and completeness validation of genome assembly.

## Code availability

No custom computer codes were generated in this work.

## Acknowledgements

This work was supported by Guangzhou Science and Technology Project [No. 201803020017], National Natural Science Foundation of China [No. 91858208], [No. 31902427], R&D Project for Jinwan Yellowfin Seabream Breeding System Construction [No. K20-42000-018], Science and Technology Project of Zhanjiang [No. 2019A03011], and Innovation Group Project of Southern Marine Science and Engineering Guangdong Laboratory (Zhuhai) [No. 311020005].

## Author Contributions

J. L. conceived the study. D. G., W. F. and J. X. collected the samples. D. G. and G. L. extracted the genomic DNA. D. G., Y. S., J. T., J. C. and K. H. assembled and curated the genome. D. G. and J. L. wrote, reviewed and edited the manuscript.

